# Mapping the drivers of within-host pathogen evolution using massive data sets

**DOI:** 10.1101/155242

**Authors:** Duncan S. Palmer, Isaac Turner, Sarah Fidler, John Frater, Philip Goulder, Dominique Goedhals, Kuan-Hsiang Gary Huang, Annette Oxenius, Rodney Phillips, Roger Shapiro, Cloete van Vuuren, Angela R. McLean, Gil McVean

## Abstract

Differences among hosts, resulting from genetic variation in the immune system or heterogeneity in drug treatment, can impact within-host pathogen evolution. Identifying such interactions can potentially be achieved through genetic association studies. However, extensive and correlated genetic population structure in hosts and pathogens presents a substantial risk of confounding analyses. Moreover, the multiple testing burden of interaction scanning can potentially limit power. To address these problems, we have developed a Bayesian approach for detecting host influences on pathogen evolution that makes use of vast existing data sets of pathogen diversity to improve power and control for stratification. The approach models key processes, including recombination and selection, and identifies regions of the pathogen genome affected by host factors. Using simulations and empirical analysis of drug-induced selection on the HIV-1 genome we demonstrate the power of the method to recover known associations and show greatly improved precision-recall characteristics compared to other approaches. We build a high-resolution map of HLA-induced selection in the HIV-1 genome, identifying novel epitope-allele combinations.

## Introduction

Variation in multiple host factors, both genetic and non-genetic, can influence the genetic composition of infecting pathogens and their subsequent evolutionary trajectory within a host. Examples include human leukocyte antigen (HLA) restriction of epitopes and subsequent escape in HIV-1 and other viruses [1-6], drug-induced selection pressure and appearance of drug-resistance mutations in viruses, bacteria and eukaryotic pathogens [7-13] and interactions between polymorphic red-blood cell types and malarial disease [14,15]. Consequently, the molecular mechanisms underlying diverse pathogen-related processes including infection, invasion, immune-response and drug resistance, can potentially be uncovered by studying the association between host factors and the genetic composition of pathogens [16-22],

However, while it is now feasible to collect large-scale data on pathogen genomic variation and host parameters, reliable hypothesis-free detection of biologically-meaningful associations between host factors and pathogen diversity is challenging for several reasons; the greatest of which is population structure. Host genetic variation often has a strong spatial structure arising from historical patterns of isolation and gene flow. Because of their commensal nature and mode of transmission, most pathogens are likely to share some of this structure, leading to non-causal association between host and pathogen genetic variation. For the same reason, geographical heterogeneity among host factors that influence pathogens causally (e.g. local variation in treatment protocols) may also lead to indirect correlation between host and pathogen genetics.

A second major challenge is statistical power. Consider searching genome-wide for associations between host genomic factors and pathogen genomic factors, in which the number of tests carried out could be in the billions. Naïve correction for multiple testing is likely to eliminate power for anything except the strongest associations. Consequently, there is a need for approaches to association testing that enable prior information about the likely structure of association to be used in the search for signal.

To date, various approaches to testing for association between pathogen genetic variation and host factors, both genetic, as in the case of classical HLA loci, and non-genetic, as in the case of drug resistance, have been developed [16,18, 23-25]. Standard association tests, which suffered the problems of stratification described above, were superseded by methods that utilise an inferred phylogeny of pathogen samples to correct for relatedness [22, 23, 26-29]. Moreover, additional power can be obtained by explicit modelling of the processes of escape and reversion in the context of HLA restriction of HIV-1 [21, 23, 24], However, such approaches have a number of limitations. For example, they do not consider recombination in pathogen genomes, they do not make use of all the data available, they typically do not infer the strength of host-induced selection or combine information across nearby sites in the context of epitope mapping and the more sophisticated approaches are often computationally prohibitive for very large samples.

To address these limitations we have developed a model-based approach to inferring the effect of host factors on pathogen genomic variation. The approach is motivated by the presence of extremely large databases of pathogen genomic data. Given the size of the databases (e.g. the Los Alamos database on HIV-1 has over 150,000 sequences encoding a portion of reverse transcriptase), it is likely that there are sequences closely related to those that infected the individuals within a particular study in question. Our approach aims to infer the most likely ancestral infecting sequence (which may be a mosaic of those in the database) for each individual and therefore to identify the pathogen evolution that has resulted directly from exposure to the current host. Moreover, we model correlations in evolution among adjacent sites within the pathogen that arise through being a shared target of selection (e.g. within a restricted HLA epitope or within a region of a protein where drug resistance and consequent compensatory mutations can evolve). We show that the approach has substantially improved power to detect sites under selection and, through applications to the evolution of drug resistance and escape from HLA-drive immune-restriction in HIV-1, how the method can deliver new insight into important biological processes.

## Method overview

Our goal is to estimate the location and nature of host-associated selection upon the pathogen genome sequence through statistical analysis of association between host factor and pathogen genomic variation. We desire our method to have four properties: the ability to use all available pathogen sequence data, irrespective of whether host factors have been measured; the ability to measure the evidence for selection across the genome analysed in a hypothesis-free manner; the ability to combine information across neighbouring positions where appropriate; and the ability to account for confounding factors such as recombination and population structure. To meet these requirements we have developed a Bayesian model-based approach in which we use an approximation to the coalescent with recombination and codon-level selection [30]. In extension to the earlier work, we enable host-specific factors to influence patterns of variation and consider the case of two data sets. The first, ***D***, represents the collection of viral sequences for which host factor data (e.g. HLA genotype or drug treatment) is also available. The second, ***D_B_***, represents a much larger dataset of viral sequences for which host factor data is not available. We use ***D_B_*** to model the set of potential viral sequences that a host can be infected with (allowing for recombination) and assume that genetic differences between this reference panel and ***D***, reflects evolution within the host. Thus host-induced selection will result in an association between host factors and the evolutionary changes observed in ***D***. A cartoon of the underlying process and our model is shown in Figure 1.

**Figure 1.**
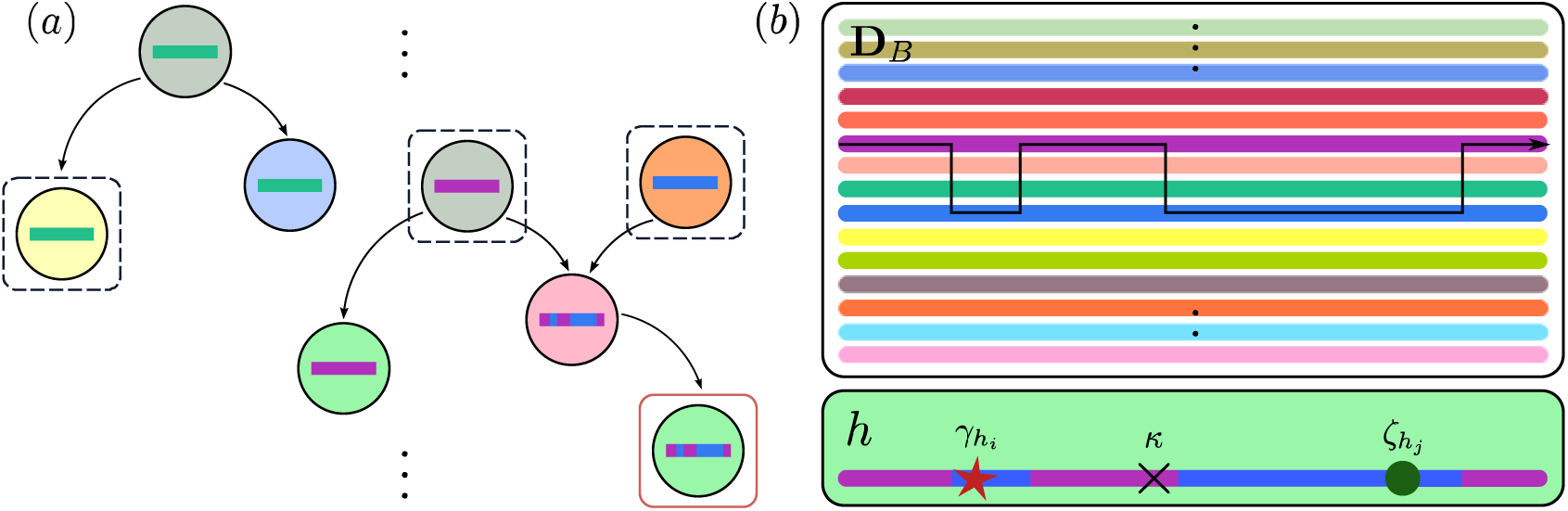
The underlying process and our approximation. *(a)* The underlying process: Infected hosts are circles, coloured by host factor information. Coloured strips represent viral consensus within infected hosts, and arrows indicate the direction of infection. Sampled hosts with pathogen sequence information are outlined. A red outline indicates that host factor information is also available, *(b)* Our approximation: Pathogen sequences in ***D*** are generated from ***D_B_***, modulated by host factors of ***D***. As |***D_B_***| » 0, we assume ***D_B_*** is the set of all possible sequences an individual can be infected with. Members of ***D*** arise from ***D_B_*** through recombination and mutation. We assume all selection along a lineage connecting each member of ***D*** to its closest neighbour in ***D_B_*** (subject to recombination) occurred within the host that the member of ***D*** was isolated from. Thus only host factor information for ***D*** is required. In *(b)*, there is one green host factor, *h.* Recombination, shown by the arrow, results in the colouring of the sequence. Three mutations have occurred: one due to *h-* associated selection (red star), one reversion (green circle), and one synonymous transversion (black cross).

The justification of the approach is that if |***D_B_***| » |***D***| then members of ***D*** will typically coalesce very recently and approximately independently with members of ***D_B_***. Thus the prior on ***D*** can be modelled with the Li and Stephens imperfect mosaic hidden Markov model (HMM), utilising a modified NY98 codon model [31]. To incorporate host induced selection, we define a ‘consensus’ codon 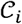 at site *i* and model selection as a scaled increase in the rate of non-synonymous change away from 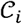. Conversely, we model reversion as a scaled increase in the codon substitution rate towards 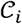. To obtain emission probabilities we integrate over the distribution of coalescent time between a member of ***D*** and ***D_B_*** assuming a coalescent model.

The HMM formulation enables efficient computation of the probability of observing a given sequence in of ***D*** given current parameter values. The product over all members of ***D*** is used to approximate the joint probability 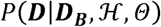, where 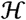 is host factor data for members of ***D*** and *θ* represents the parameters of recombination and codon substitution. To speed up computation, we *a priori* identify a subset of ***D_B_*** for each member of ***D*** that is used to model its ancestor. The model parameters we aim to infer are: the synonymous transition rate, *μ*, the *dN/dS* ratio at each site, *ω_i_*, the recombination probabilities between neighbouring sites, *r_i_*, ithe rate of reversion *ζ_i_* and the host-factor-dependent scaling of escape rate at each site, *γ_h,i_*. Other parameters (transition-transversion rate and fraction of codon substitutions more than one nucleotide change away) are estimated from external data. We use Markov chain Monte Carlo to sample from the posterior distribution for all parameters. Full details of the methodology are given in the Methods and Supplementary Methods: Supplementary Text 1.

## Simulations and methods comparison

To validate our method and compare its performance to alternative approaches for detecting host-induced selection we carried out two simulation studies. First, we simulated data under the fitted model to evaluate power and accuracy. We used HIV-1 protease sequence data from three major public databases as a reference (Table 1; *n* = 146,523), and simulated 100 replicates of a data set of 460 study sequences with six different HLA-induced selection profiles, four of which were common (n = 100 each) and two of which were rare (n = 30 each); Supplementary Figure 1. We performed simulations with recombination ranging between 0 and 0.01 (a rate of 0.01 meaning that on average the ancestor of each study sample copies 100 contiguous codons from a reference sample before recombination) and with the size of the selected sample-specific ancestral reference panel ranging from 10 to 100. In the absence of recombination, parameter estimates are largely unbiased and accurate, although the rate of reversion has high variance (Figure 2A; Supplementary Figure 2). Estimates for rare alleles perform similarly to those for common alleles (Figure 2B). We find little impact of the size of the sample-specific reference data set and the posterior distributions are well calibrated (Supplementary Figure 3). Recombination leads to a downward bias in estimates of recombination and selection intensity and upward bias in the rate of reversion (Figure 2C; Supplementary Figure 4) and consequently poorer posterior calibration at sites under selection (Supplementary Figure 4). Nevertheless, the inferred profiles of host-dependent and host-independent selection pressures remain strongly correlated to the truth. We also considered the impact of error in choosing the sample-specific reference panel by forcing inclusion of the sequences actually copied. We find that forcing the true ancestors to be included improves accuracy by only a small margin (Supplementary Figure 5) indicating that 100 potential ancestors chosen through Hamming distance is sufficient.

**Figure 2.**
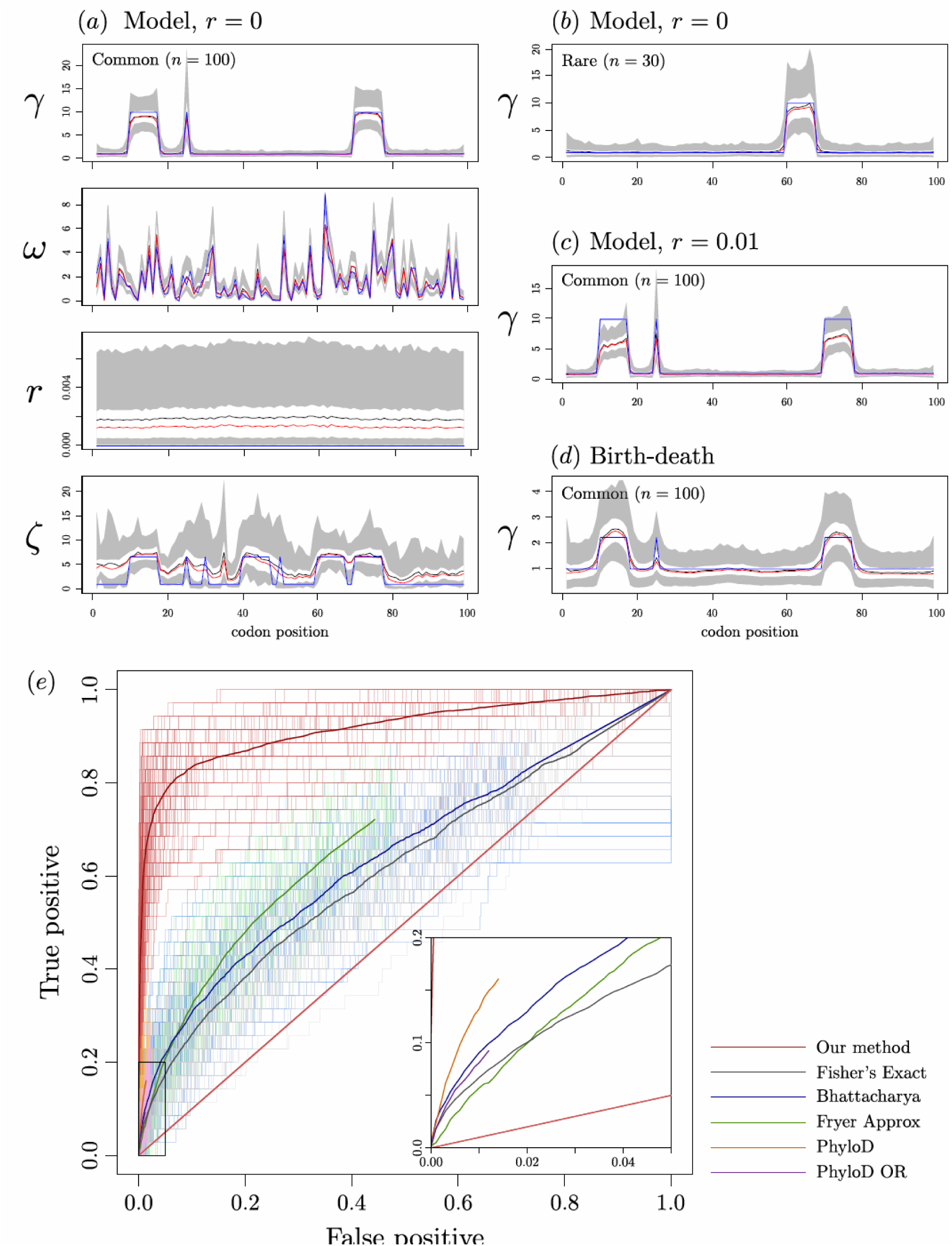
Simulation results summary. Inference results for Simulation studies 1 and 2 are shown in (*a-d*). (*a*) Simulation study 1; *r* = 0. *γ* for a common HLA (*n =* 100), *dN / dS* ratio, recombination probability between adjacent sites (*r*), and reversion scaling (ζ). *(b)* Simulation study 1; *r* = 0, *γ* for a rare HLA (*n =* 30). *(c)* Simulation study 1; *r* = 0.01, *γ* for a common HLA *(n =* 100). *(d)* Simulation study 2, *γ* for a common HLA *(n =* 100). In *(a-d)* (except *ω* in (*a*)), averages are taken over 100 independent MCMC runs on independent simulated data with the same underlying parameters. The true value is shown in blue, mean and median estimates are in black and red respectively. White bands enclose 50% credible intervals, in turn enclosed by grey 95% credible intervals. In *(c)*, the truth is rescaled by the average number of individuals between a randomly chosen pair of leaves in the sampled birth-death tree. For *ω* in *(a)*, averages are taken over a single MCMC run, chosen at random as *ω* differs across simulation runs (sampled from the prior). See Figures S2-S5 for full results summaries, *(e)* ROC curves for existing method. Six methods used to identify HLA associated selection on viral sequence are applied to data simulated under the birth-death process used in simulation study 2. The Inset zooms in to the region enclosed by the black box. ROC curves for 100 independent birth-death simulations are lightly coloured, and averaged to generate the heavier lines. ROC curves for Fryer Approx, PhyloD and PhyloD OR do not extend to (1,1). For Fryer Approx this is because we stop our threshold for estimated rates at 0. For PhyloD and PhyloD OR, it is because many sites will not be included in leaf distribution or logistic regression respectively.

**Table 1.** Data used in this study.

| Database | Cohort size | Details |  |  |  |
| --- | --- | --- | --- | --- | --- |
| Los Alamos HIV Sequence Database [44] | Protease $n = 119,878$<br>Reverse transcriptase $n = 148,866$ | Fed by biweekly downloads of new HIV sequence data submitted to GenBank. Metadata is extracted from GenBank submissions and manually annotated with additional information taken from the corresponding publications and through direct interaction with authors [45]. | | | |
| Stanford Drug Resistance Database [9] | Protease $n = 81,533$<br>Reverse transcriptase $n = 88,780$ | Collection of protease and reverse transcriptase sequences with associated patient drug prescription data. | | | |
| HIV positive selection database [46] | Protease $n = 45,161$<br>Reverse transcriptase $n = 45,161$ | Protease and reverse transcriptase sequence data taken from HIV-1 patient plasma samples by Specialty Laboratories from 1999 to 2002 [46]. The database was created in order to identify regions of drug selection using estimates of non-synonymous/synonymous base changes along the viral genome. | | | |

| Dataset | Cohort size | Sampling date | Geographic region | Treatment | Study requirements |
| --- | --- | --- | --- | --- | --- |
| SSI/TT [47, 48] (Swiss portion) | $n = 79$ | 2000 | Various cities across Switzerland | HAART | Undetectable VL for $> 6$ months, $CD4^+$ count $> 300 \mu l^{-1}$. No history of non-nucleoside reverse transcriptase inhibitors. |
| SPARTAC [44] (UK portion) | $n = 258$ | Aug 2003 – July 2007 | UK | ART naïve on recruitment | Primary infection, though definition of this is complex, see [44] for details. |
| Bloemfontein [49, 50] | $n = 278$ | Feb 2006 – Sept 2006 | Bloemfontein, South Africa | ART naïve | Chronic infection, low and high $CD4^+$ count favoured: 96 high ($> 500 \mu l^{-1}$), 18 medium ($200 - 400 \mu l^{-1}$), 164 low ($< 100 \mu l^{-1}$). |
| Durban [51, 52] | $n = 1218$ | 1999-2006 | Durban, South Africa | ART naïve | Recruited following voluntary counselling and testing in antenatal or outpatient clinics. |
| Mma Bana [53] | $n = 514$ | July 2006 – May 2008 | Gaborone, Botswana | ART naïve | Pregnant women. |
| Los Alamos | Protease $n = 432$<br>Reverse transcriptase $n = 334$ | All remaining sequences with associated host HLA data available in the Los Alamos HIV sequence database [54] which were not present in the above studies. | | | |

In the second set of simulations we assessed robustness by simulating data under a birth-death model (without recombination), setting the current infected population size at 1,000,000 and sampling proportion at 10% to define ***D_B_*** (Supplementary Text 2). We find some attenuation of the selection signal (and over-estimation of the reversion parameter) but strong correlation between the inferred and estimated strengths of host-dependent and host-independent selection (Figure 2C; Supplementary Figure 6).

The birth-death simulations also enables comparison of our approach with five alternative methods for identifying sites under host-factor specific selection (Supplementary Text 3): Fisher’s exact test (as in Moore *et al.* [16]); a phylogenetically corrected Fisher’s exact test (as in Bhattacharya *et al.* [22]); an approximate escape rate estimate (as in Fryer *et al.* [18]); a ‘Phylogenetic dependency networks’ approach - PhyloD (as in Carlson *et al.* [26]); and PhyloD OR (as in Carlson *et al* [21]). Note that the phylogenetic approaches can only be applied if there is no recombination. In each case, we assume that the ‘wild-type’ strain is correctly defined and set any non-synonymous difference from consensus as a candidate for escape. For each method, we obtained p-values or parameter estimates that provide a metric for the strength of selection conditional on each HLA type and generated receiver operating characteristic (ROC) curves for each simulation (Figure 2E). We find that our approach dramatically increases sensitivity for a given false positive rate (FPR). For example at FPR = 0.01, our sensitivity is 0.65, compared to the second best performing method, PhyloD, at 0.13. We therefore conclude that augmenting study data with a large reference data set and modelling the escape process explicitly provides a substantial gain in the ability to identify sites under host-factor specific selection.

## Drug associated selection

To provide empirical validation of our approach for detecting host-factor dependent selection we analysed HIV-1 data from the Stanford drug resistance database [9] in which viral sequences are linked to antiretroviral drug treatment history of the patient. We therefore aim to learn drug-treatment specific evolution, while other selective factors (e.g. selection due to CTL pressure, the antibody response, or the APOBEC3G response) will be captured by *ω.* We analysed protease and reverse transcriptase independently and excluded integrase due to lack of data [9]. Priors on parameters are given in Supplementary Table 1. We defined the collection of sequences in hosts not receiving any therapy at the time of sequencing as ***D_B_***, and viral sequences in hosts receiving any treatment (coupled with their drug regime data) as ***D***. We then randomly assign 2000 sequences from ***D_B_*** to ***D***. This ensures that drug-associated selection common to all treatments is not absorbed into *ω*. Full details of data preparation are given in Supplementary Text 4.

We estimated *γ_i_*_,*h*_ for protease and reverse transcriptase inhibitors prescribed to at least 10 individuals in ***D*** (Figure 3A; Supplementary Figure 7) and identified sites with very strong evidence for drug-induced escape (median estimate of selection factor > 2 and at least 97.5% of the posterior > 1; top-tier) and moderate evidence (median estimate of selection factor > 1.5 and at least 90% of the posterior > 1; second tier). We compared the enrichment of sites identified to known major drug resistance mutations (DRMs) [9] (Figure 3B). For some drugs, DRM data was lacking because the drug is no longer commonly used (DDC and DLV), was an experimental drug that failed (*α*APA, and ADV) or is now used in small amounts with other drugs (RTV). We observed strong, statistically significant and consistent enrichment of DRMs at sites identified as selected; with elevated enrichment (as measured by odds-ratio) in the strongest DRMs and sites with strongest evidence for selection. For example, of the 31 strongest DRMs in RT, 11 are found in the top tier selected sites and a further 6 in the second tier. Of the 41 apparent false positive sites, 17 are described as being selected for in the literature (but not classified as a major drug resistance mutation in the Stanford database); for example mutation at codon 11 of RT is known to be associated with minor reductions in APV susceptibility [32], but is not classified as a major DRM [9], A further three apparent false positive are sites of DRMs for treatment cocktails including the drug, nine are sites of DRMs for drugs in the same class but where more specific information is not provided, seven are sites of DRMs for different drugs in the same class and only five appear to have no support in the literature (Supplementary Table 2). False negatives are caused by sites where the selected codon is more than one nucleotide change away from the consensus (five mutations), one is an insertion, eight have *in vitro* support for drug resistance but are not documented as being selected for *in vivo*, seven have no compelling evidence in the literature and three are genuinely missed (Supplementary Table 3) [9]. Further evidence for the biological validity of the inferred profiles is given by the co-clustering of related drugs in trees constructed from the selection intensity profiles (Supplementary Figure 8). In summary, we find that the method can identify sites selected by specific drug treatments *in vivo*, either directly through resistance or indirectly through compensation for resistance mutations that reduce fitness.

**Figure 3.**
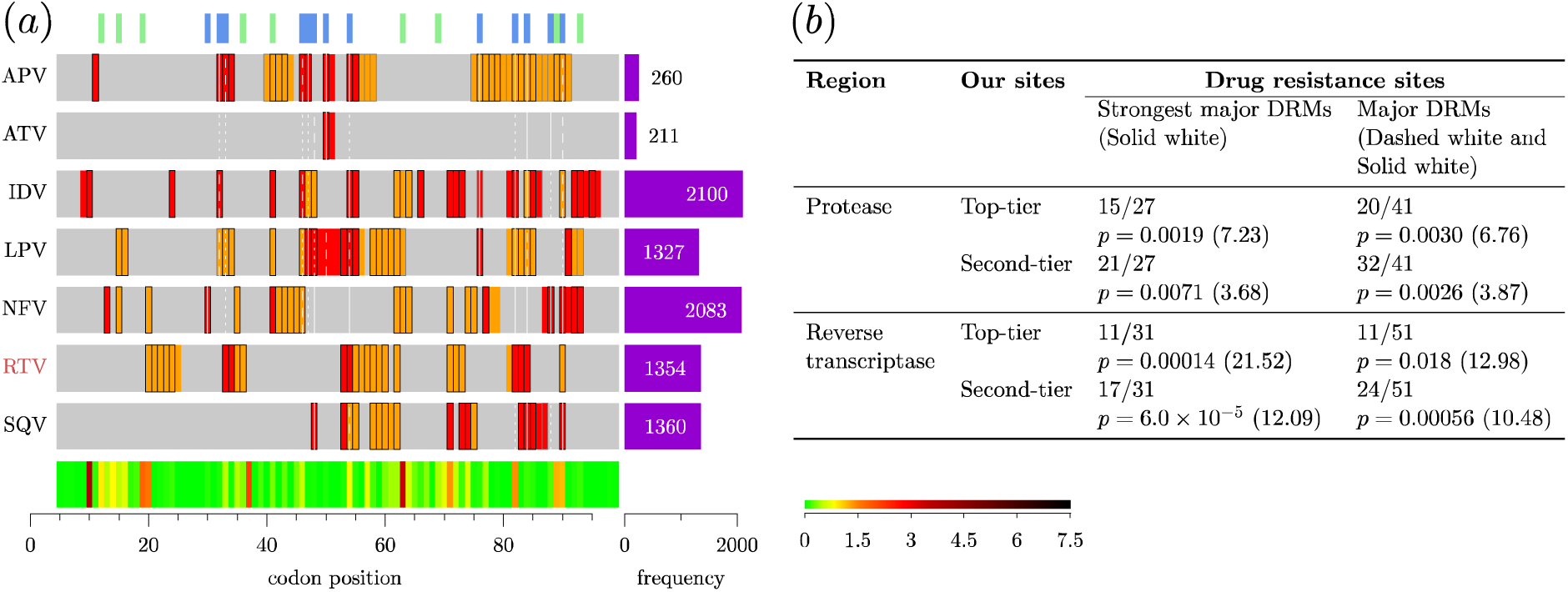
Drug associated selection analysis. *(a)* Protease results summary. Grey bars highlight the region analysed. Codon position is measured relative to the HXB2 from the start of protease. Rows summarise drug-associated selection - drugs are shown to the left. Purple bars show the number of individuals prescribed the drug at viral sampling. Red and orange indicate our median estimate is > 2 and > 1.5 respectively. Sites are outlined in Black and grey if the 2.5% and 10% quantile is > 1 respectively. Green and blue lines at the top of the plots highlight differences between subtype B/C at the amino-acid level, and sites of DRMs [9] respectively. Classes of DRM are displayed: solid white lines tag sites of DRMs which ‘confer the highest levels of resistance’ for that drug, dashed white lines tag ‘major’ DRMs, and dotted white lines tag DRMs ‘when combined with other mutations’ [9]. Drugs highlighted red do not have major DRMs in the Stanford drug resistance database. Median *ω* is displayed at the foot of *(a)*, according to the colourbar. Key: APV = Amprenavir; ATV = Atazanavir; IDV = Indinavir; LPV = Lopinavir; NFV = Nelfinavir; RTV = Ritonavir; SQV = Saquinavir, *(b)* Sensitivity of our inference. Proportions of the strongest major DRMs and major DRMs we identify when using our top-tier and second-tier candidate sites are provided, with associated p-values and odds-ratios (OR).

## HLA-associated selection

While the extent to which host HLA alleles can influence patterns of escape and reversion in HIV-1 has been studied extensively, the methods developed here provide additional power and accuracy as well as provide control against factors such as population stratification and recombination. Moreover, they also provide a framework in which to combine data from multiple previous studies. We have assembled nearly 3,000 HIV-1 sequences from patients with known HLA class I genotypes from six studies, of European and African ancestry representing a mixture of subtype B and C sequences (Table 1). Moreover, we augment the data with HIV-1 genome data taken from three sources, representing 250,000 − 280,000 sequences depending on the gene (Table 1). We analysed the data in two ways, first by considering escape as deviation from subtype B and also be considering escape from subtype C. We expect these analyses to yield similar results, except at sites where subtypes differ systematically. We analysed data from protease and reverse transcriptase and considered all HLA loci jointly. Priors on parameters are given in Supplementary Table 1. Full details of data preparation are given in Supplementary Text 4. We report results for all alleles for which we have over 10 copies. We define ‘top-tier’ HLA associated candidate sites as those where the median *γ_i,h_* > 2 and the lower 2.5% quantile > 1, and ‘second-tier’ candidate sites as those where the 10% quantile of *γ_i,h_* > 1.

To illustrate the validity of our approach, we first considered the B*51 restricted epitope TAFTIPSI in reverse transcriptase (Figure 4). We find a strong signal of selection within the epitope, with the escape site (position 135) experiencing the strongest rate elevation. Interestingly a variant at site 129 which is known to abrogate CTL recognition *in vitro* [33], is highly conserved *in vivo* and shows no signal for selection, presumably due to other fitness consequences. In contrast, we also see strong evidence for B*51 associated escape around codons 173 and 195, which have not been reported previously. Other known allele-epitope combinations that we recover include, for example, B*18 associated selection around codon 138 [34-36].

**Figure 4.**
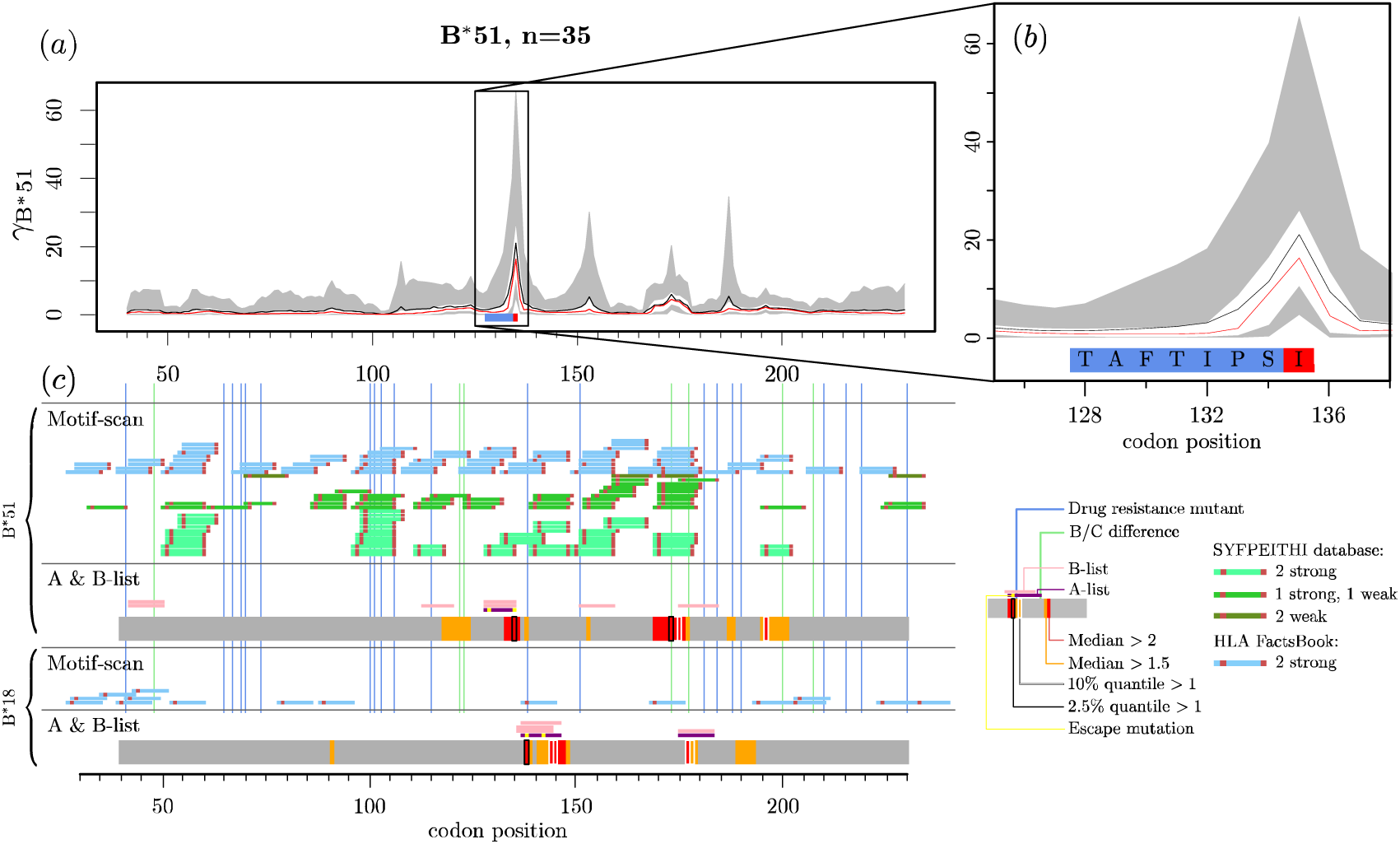
Selected results summaries. *(a) B*\*51 associated selection away from 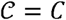across the analysed region of reverse transcriptase. Site numbering is relative to the HXB2, starting from beginning of reverse transcriptase. Plotting is as in Figure 2 *(a-d).* The *B*\*51 epitope TAFTIPSI is highlighted by the blue box – the site of the known escape variant is highlighted red. The black rectangle is zoomed in on in *(b). (c)* Epitopes predicted by Motif Scan and A-list/B-list epitopes for two example HLA types are displayed. Corresponding results from our inference are shown on the grey strips – colouring is summarised by the key.

To compare to experimental and previous work on epitope restriction we classify documented epitopes into an A-list, which represents the best-defined experimentally determined HIV-1 CTL epitopes, updated yearly [13] and a B-list, which refers to the entire collection of epitopes reported in the literature [37], Among the A-list epitopes, some have documented CTL escape variants, either *in vivo* or *in vitro.* See Supplementary Table 4-5 for details and Supplementary Text 5 for discussion of each case. We also consider an *in silico* set of predictions for ‘strongly-binding’ anchor residues generated by Motif-Scan [14]. We compare these sites to our estimates of HLA-associated selection across protease and reverse transcriptase (Figure 5, Supplementary Figures 9-13).

**Figure 5.**
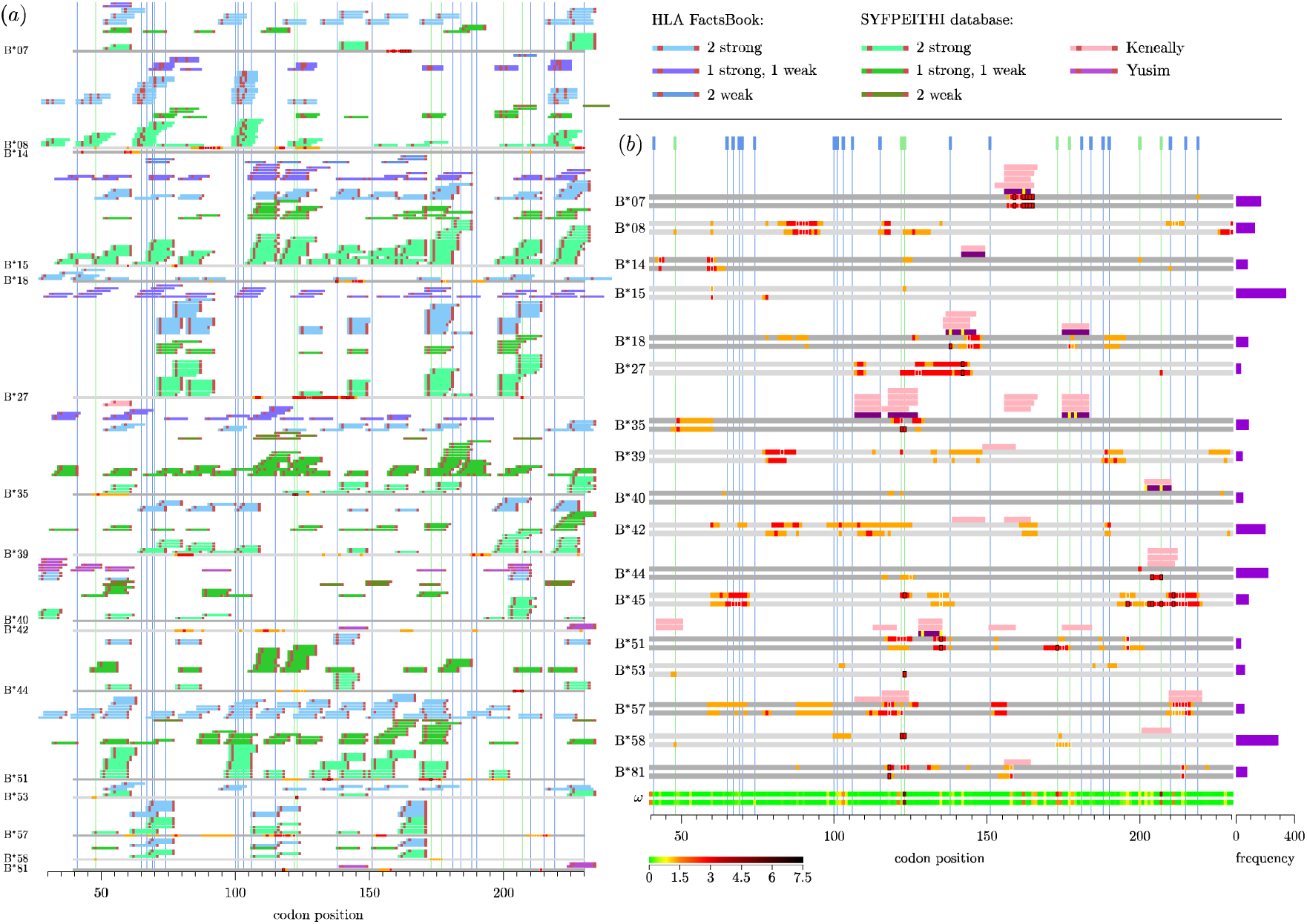
HLA-B associated selection in reverse transcriptase. *(a)* Motif scan compared to our inference. Inference of HLA associated selection from 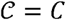 is summarised on the grey strips: sites with median *γ_i,h_ >* 2 and 1.5 are coloured red and orange respectively, sites with 2.5% and 10% quantile > 1 are outlined in black and white respectively. Blue and green vertical lines tag sites of drug resistance mutations and sites that differentiate subtype B/C viruses at the amino acid level. We scan reverse transcriptase for putative epitopes matching to known binding motifs in the literature; see key. Anchor residues are highlighted in red. *(b)* A-list/B-list compared to our inference. Inference results on the grey strips are coloured as in *(a).* Two grey strips per HLA show selection away from 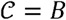and 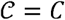 respectively. Bar plots to the right show HLA frequencies. The two lowest strips summarise median *ω*, see colour bar. A-list [43] and B-list [37] epitopes above the grey are coloured purple and pink respectively. Sites of known escape variants within the A-list epitopes are highlighted yellow.

To assess overlap between sites identified we measured the enrichment of sites identified as under selection and used a permutation strategy to assess significance (Supplementary Text 7). Across protease and reverse transcriptase we find strong evidence for enrichment of top-tier signals of selection at A-list epitopes for HLA-B (Table 3). As the confidence in the epitope collection or the strength of selection decreases, so does the enrichment. We find no evidence for enrichment of selection at computationally predicted epitopes and little evidence for overlap of HLA-A associated sites, though in general we note that HLA-A alleles show much weaker evidence for selection in general (Supplementary Figures 9 and 12). We also find very little evidence for HLA-C driven selection at all (Supplementary Figures 10 and 13).

**Table 2.** Overlap between sites under selection and known HLA epitopies

| Region | HLA | Selected sites | Epitope collection: p-value (odds-ratio) |  |  |
| --- | --- | --- | --- | --- | --- |
|  |  |  | Motif scan<sup>a</sup> | B-list | A-list |
| Protease | A | Top-tier | 0.693 | 1 (0) | 1 (0) |
|  |  | Second-tier | 0.0855 | 0.160 (4.12) | 0.166 (9.00) |
| | B | Top-tier | 1 | 0.0420 (48.8) | 0.0499 (∞) |
|  |  | Second-tier | 0.964 | 0.213 (3.50) | 0.0335 (14.0) |
| Reverse transcriptase | A | Top-tier | 0.953 | 0.404 (1.28) | 1 (0) |
|  |  | Second-tier | 0.935 | 0.122 (1.57) | 0.499 (1.61) |
|  | B | Top-tier | 0.137 | 0.0000290 (55.2) | 0.00278 (63.3) |
|  |  | Second-tier | 0.303 | 0.000250 (19.4) | 0.00278 (37.7) |

<sup>a</sup>Putative epitopes were generated using the subtype C consensus; results using the subtype B consensus were similar.

**Table 3.** Codons in reverse transcriptase showing evidence for HLA associated selection

| HLA | $C = B$ | $C = C$ |
| --- | --- | --- |
| A*03 | 135, 166† | 135, 136 |
| A*11 |  | 126 |
| A*43 | 118, 135, 139, 140, 141, 142 | 118, 122*, 123*, 135 |
| A*66 | 135 | 135,<u>138</u> |
| B*07 | 159, 162†, 163, 164, 165 | 159, 162†, 163, 164, 165 |
| B*18 |  | <u>138</u>† |
| B*27 | 142 | 142 |
| B*35 |  | 122*, 123* |
| B*44 |  | 204, 207* |
| B*45 | 123*, 211 | 196, 203, 204, 207*, 211 |
| B*51 | 135† | 135†, 173* |
| B*53 |  | 123* |
| B*58 | 122*, 123* |  |
| B*81 | 118 | 118 |
| C*04 | <u>210</u>, 211 |  |

Sites defined by $\gamma_{i,h} > 2$ and lower 2.5% quantile $> 1$ . Underline: Site of known drug resistance mutations. \*: Site differentiates subtype B and C amino-acid consensus. †: Known HLA-site association.

Overall, only a small number of sites identified here as being HLA associated have been described previously; four in reverse transcriptase (Table 3),and one in protease (Table 4). Conversely, not all previously reported escape mutation sites are identified here. In some cases this may be due to low sample size in this study. However, it is also possible that earlier studies failed to account adequately for linkage disequilibrium between HLA alleles. We also identified potentially novel signals of HLA-driven epitope escape. For example, B*45 shows multiple signals of selection around codon 200 of reverse transcriptase (Figure 5), yet there are no reported epitopes. This may reflect a historical bias towards studies being carried out in European-ancestry populations with subtype B viruses where some HLA alleles and viral epitopes are rare. However, we also note that results between the subtype-B and subtype-C consensus analyses are highly concordant, with a few interesting exceptions. For example, B*58 shows evidence for inducing strong selection away from the subtype B consensus around codon 122-123 of reverse transcriptase but not away from the subtype C consensus. Interestingly, this is a position of divergence between subtype B and subtype C viruses and B*58 is typically more common in populations with HIV-1 subtype C viruses, suggesting that B*58-induced selection pressure may have driven the fixation of the difference between subtypes.

**Table 4.** Codons in protease showing evidence for HLA associated selection

| HLA | $\mathcal{C} = B$ | $\mathcal{C} = C$ |
| --- | --- | --- |
| A*31 |  | 57, 61, 62, 64 |
| A*66 | 35, 36*, 37 |  |
| B*13 |  | 62, 63*, 64 |
| B*44 | 35†, 37, 38, 39 | 35†, 36, 37, 38, 39, 41* |
| B*49 | 64, 65, 66, 67, 68 | 61, 62, 63*, 64, 65, 67, 68 |
| B*51 |  | 37 |
| C*08 | 14 |  |
| C*18 | 35 | 15*, 35 |

Sites with $\gamma_{i,h} > 2$ and lower 2.5% quantile $> 1$ . Underline: Site of drug resistance mutations. \*: Site differentiates subtype B and C amino-acid consensus. †: Known HLA-site association.

To assess the relationship between sequence similarity between HLA alleles and the inferred HLA-induced selection profiles, we measured the concordance between dendrograms inferred from pairwise differences of classical HLA alleles protein sequences and dendrograms inferred from the estimated selection profiles (Supplementary Text 8). We find that closely related HLA alleles have closely related selection profiles for HLA-A (odds-ratio = 2.1; permutation p-value = 0.017) and HLA-B (odds-ratio = 3.1; permutation p-value = 0.014), but not HLA-C (odds-ratio = 0.71; permutation p-value = 0.71). However, deeper structure within the inferred trees shows little concordance.

Finally, as noted recently [38], a number of sites both show evidence of HLA-driven selection as well as being associated with evolution in response to drug treatment (for example, around codon 70 in reverse transcriptase for B*45). Depending on whether the two types of selection act in the same or different directions, such pressures could either speed up or delay the origin of drug resistance. These results suggest that HLA genotype could be a predictor of the response to drug treatment.

## Discussion

Differences between hosts, such as in their immune system, or treatment received, can lead to host-specific selection regimes and subsequent adaptation by the pathogen. If within-host adaptation is at a cost to intrinsic fitness, then, on further transmission, selection may favour reversion to the fitter, ancestral state. To learn about such forces, the ideal experiment is to compare the genetic composition of the original infecting pathogen strain and that present sometime after successful infection across a large number of hosts that differ only in the factors of interest. While only strictly possible in experimental systems, observational approaches aim to learn about the same processes, but, to do so, have to make assumptions about the (unobserved) pool of pathogens to which infected individuals were exposed and the distribution of potential confounding factors. If assumptions are not met, power to detect true associations will be reduced and there is a risk of false positive results.

The introduction of phylogenetic methods for association testing in pathogen genomics [20-23, 26, 27, 39] provided a partial solution to many issues, by using genome-wide relatedness between pathogens as a proxy for correlation in unobserved confounding (such as population structure), similar to the use of principal components in the analysis of human genetic association [40]. Moreover, by developing explicit models of molecular evolution in response to host factors, it is possible to learn relative or even actual rates of escape and reversion [23]. However, phylogenetic methods have limitations. For example, they can be computationally expensive, meaning they are hard to apply to huge data sets; assumptions of homogeneity in rates over time and space typically have to be made; and, more fundamentally, recombination is widespread among pathogens.

To address these shortcomings, we have developed a model-based approach to association testing that exploits the availability of extremely large reference data sets on pathogen variation (but where there is no relevant metadata concerning the factors of interest), which are likely to contain genomes closely related to the samples of interest (with relevant metadata). Moreover, we argue that when identifying host-driven selection, most of the information lies near the tips of the trees as deeper comparisons have to integrate over many transmission events. By utilising the Li and Stephens [41] haplotype model, combined with previous work on modelling adaptive evolution and reversion [18, 23, 30] we have developed an approach that both dramatically increases power to detect associations and scales to huge data sets.

The analysis of *in vivo* patterns of evolution provides a complementary approach to the *in vitro* study of drug resistance and immune evasion. The evolutionary response reflects the combination of fitness gains of escape and the fitness loss through modification. Hence sites with strong resistance may have too strong a fitness cost to be typically selected for and weak resistance changes that have little fitness impact may be strongly selected for. Moreover, compensatory changes in the pathogen genome can potentially counteract fitness loss and hence appear as part of the resistance response. Our analysis of drug-associated evolution using data from the Los Alamos database identifies the majority of major DRMs and sites not identified typically result from failings of the model (variants that are more than one change away from the consensus codon, insertions and deletions). However, we also identify novel sites with compelling evidence for drug-associated selection, likely reflecting minor DRMs with small fitness loss and compensatory changes.

The analysis of HLA-associated selection presented here is the most complete and comprehensive to date, covering six studies and nearly 3,000 HIV-1 sequences with linked patient HLA class I data. We show the method can recover well-known epitopes and associated escape mutations and that there is strong overlap between curated lists of epitopes and those identified here. However, we also found that simple *in silico* epitope prediction fails to provide a good predictor of the HIV-1 evolutionary response and multiple cases where curated epitopes do not appear to lead to a substantial selective pressure. We also showed a preponderance of signals of escape associated with HLA-B alleles, as opposed to HLA-A and HLA-C and identified a number of sites where differences between subtype consensus sequences collocate with sites selected by particular HLA alleles, raising the question of whether differences in HLA allele frequency between populations could have contributed to the differentiation of HIV-1 subtypes [42].

As field-based sequencing of pathogens becomes feasible, so does the potential to accumulate vast data sets that can track spatial and temporal shifts in selection pressures affecting pathogens and their responses. Such studies will require new approaches to studying genetic association. While the method developed here is aimed at detecting within-host evolutionary responses, future extensions could include adaptation at a population level, for example through exploiting longitudinal sampling and/or spatial heterogeneity in treatment.

## Online Methods

We perform inference via Markov chain Monte-Carlo (MCMC) using the Li and Stephens approximation to the coalescent with recombination [41], with one further key approximation. Consider two datasets:

1. ***D*** *=* viral sequences for which host HLA data is also available.
2. ***D_B_*** *=* viral sequences for which host HLA data is not available.

As ***D_B_*** is very large, we can assume that ***D_B_*** represents the collection of viral sequences (subject to recombination) that a host can be infected with. A large number of sequences in ***D_B_*** results in short coalescence times between any given member of ***D*** and ***D_B_***, increasing our power to estimate parameters. By making this approximation, we may then assume that selection along the lineage joining a given member *d* of ***D*** to its nearest neighbour in ***D_B_*** occurred within the individual from whom the virus *d* was sampled. We have associated host HLA profiles for ***D***, and so may then estimate HLA-associated selection conditional on this HLA information. A cartoon of this model is shown in Figure 1b. Full details of our approach and inference regime are provided in Supplementary Text 1.

The Li and Stephens approximation allows us to evaluate the probability that a member of ***D*** arises as an imperfect mosaic of sequences in ***D_B_*** using a hidden Markov model (HMM). To model this process, we must specify a recombination model, and model of sequence evolution. We assume that recombination is site dependent, and occurs with probability *r_i_* between site *i* – 1 and *i* to any other member of ***D_B_*** uniformly at random. We model sequence change using the NY98 codon model [31] (which distinguishes transitions from transversions via the transition/transversion ratio; k, and nonsynonymous changes from synonymous changes via the *dN/dS* ratio *ω*), with some modifications. To incorporate HLA-associated selection for escape, we define a ‘consensus’ non-escaped strain 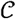 for the region to be analysed. A scaling, Π*_h∈H_γ_i,h_*, dependent on the host’s HLA profile, ***H***, then modifies any non-synonymous codon change from this consensus strain. We model reversion in a similar manner: a boost in codon substitution rate towards the consensus codon 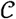 at each site, *ζ_i_*. To switch from rates to probabilities we integrate over the length of a lineage connecting a member of ***D*** to ***D_B_*** assuming a standard coalescent. For example for a non-synonymous transition from 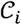 to some codon *C*,

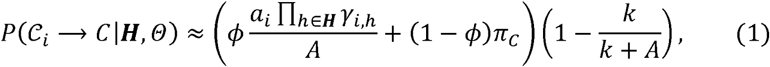

where *ϕ* is the empirically estimated probability of observing a single codon change along the lineage joining ***D*** to ***D_B_****, A* is the total codon substitution rate out of 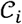, *α_i_* is the number of nonsynonymous transitions from *C_i_,k* is the number of sequences in ***D_B_*** and *π_c_* is the empirical probability of observing *C* given a ≥2 step codon change. We approximate the probability in this manner to avoid computationally expensive matrix exponentiation (Supplementary Text 1). Given recombination probabilities and codon transition probabilities for all possible codon changes and the Li and Stephens approximation, we are armed with an HMM to describe an approximation to the process generating members of ***D***. We can therefore use the forwards and backwards algorithms to integrate over all paths through ***D_B_*** to generate each member of ***D*** in turn. We then take the product over all members of ***D*** to approximate the likelihood:

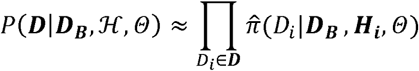

where 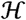 HLA information (across members of ***D*** and ***D_B_***)*, **H_i_*** is the collection of HLA types of the host associated to viral sample *D_i_* ∈ ***D***, and 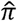 is the function describing the Li and Stephens approximation with our model of recombination and codon evolution.

Evaluating 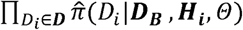 in practice becomes computationally difficult as ***D_B_*** increases in size, as it involves the evaluation of arrays of dimension |***D_B_***| × |***D***| × *|codon sequence*|. To increase computational efficiency, we restrict ***D_B_*** to a subset 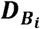 for each member of ***D***. In our simulations and analyses we use a simple Hamming distance restriction: the closest *n* sequences by Hamming distance to define each 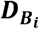.

Now that we are able to evaluate approximate likelihoods rapidly, we can perform MCMC to obtain posterior distributions for our parameters of interest:

- *dN/dS* at each site, *ω_i_*.
- H LA-associated selection at each site, *γ_h_,_i_*.
- Recombination probabilities between neighbouring sites, *r_i_*.
- Synonymous transition rate, *μ.*

Move proposals to explore that space of parameters and our inference model are provided in the Supplementary Text 1.

## Acknowledgements

Funded by an EPSRC studentship to DP, Wellcome Trust grant 100956/Z/13/Z to GM, Wellcome Trust grant WT104748MA to PJRG, National Institutes of Health grant R01AI46995 to PJRG, National Institute of Allergy and Infectious Diseases grant U01-AI066454 to RS, European Union grant SANTE/2007/147-790 to DG and CV. JF funded by the MRC. We thank Hannah Roberts for methods discussions.

## Code and data availability

Software for performing analyses is available from https://github.com/astheeggeggs/mcqueen under the MIT licence.

## References

1. Goulder, P. and B. Walker, HIV and HLA Class I: An Evolving Relationship. Immunity, 2012. 37(3): p. 426–440.

2. Woolthuis, R., et al., Long-term adaptation of the influenza A virus by escaping cytotoxic T-cell recognition. Scientific Reports, 2016. 6: p. 33334.

3. Yauch, L, et al., A Protective Role for Dengue Virus-Specific CD8+ T Cells. The Journal of Immunology, 2009. 182(8): p. 4865–4873.

4. Bowen, D. and C. Walker, Adaptive immune responses in acute and chronic hepatitis C virus infection. Nature, 2005. 436(7053): p. 946–952.

5. Phillips, R., et al., Human immunodeficiency virus genetic variation that can escape cytotoxic Tcell recognition. Nature, 1991. 354(6353): p. 453–459.

6. Butler, N., et al., Structural and Biological Basis of CTL Escape in Coronavirus-Infected Mice. The Journal of Immunology, 2008. 180(6): p. 3926–3937.

7. Qing, M., et al., Characterization of dengue virus resistance to brequinar in cell culture. Antimicrobial agents and chemotherapy, 2010. 54(9): p. 3686–3695.

8. Sandgren, A., et al., Tuberculosis Drug Resistance Mutation Database. PLOS Medicine, 2009. 6(2): p. e1000002.

9. Shafer, R.W., Rationale and uses of a public HIV drug-resistance database. J Infect Dis, 2006. 194 Suppl 1: p. S51–8.

10. Shafer, R. and J. Schapiro, HIV-1 drug resistance mutations: an updated framework for the second decade of HAART. AIDS reviews, 2008.10(2): p. 67–84.

11. Chen, Z.-W., et al., Global prevalence of pre-existing HCV variants resistant to direct-acting antiviral agents (DAAs): mining the GenBank HCV genome data. Scientific reports, 2016. 6.

12. Martinez, J.L. and F. Baquero, Mutation Frequencies and Antibiotic Resistance. Antimicrobial Agents and Chemotherapy, 2000. 44(7): p. 1771–1777.

13. Tanwar, J., et al., Multidrug Resistance: An Emerging Crisis. Interdisciplinary Perspectives on Infectious Diseases, 2014. 2014: p. 1–7.

14. Zimmerman, P., et al., Red blood cell polymorphism and susceptibility to Plasmodium vivax. Advances in parasitology, 2013. 81: p. 27–76.

15. Lell, B., et al., The role of red blood cell polymorphisms in resistance and susceptibility to malaria. Clinical infectious diseases: an official publication of the Infectious Diseases Society of America, 1999. 28(4): p. 794–799.

16. Moore, C., et al., Evidence of HIV-1 adaptation to H LA-restricted immune responses at a population level. Science (New York, N.Y.), 2002. 296(5572): p. 1439–1443.

17. Palmer, D., et al., Integrating genealogical and dynamical modelling to infer escape and reversion rates in HIV epitopes. Proceedings of the Royal Society B: Biological Sciences, 2013. 280(1762): p. 20130696.

18. Fryer, H., et al., Modelling the Evolution and Spread of HIV Immune Escape Mutants. PLoS Pathog, 2010. 6(11): p. e1001196.

19. Apps, R., et al., Influence of HLA-C Expression Level on HIV Control. Science, 2013. 340(6128): p. 87–91.

20. Carlson, J., et al., Leveraging Hierarchical Population Structure in Discrete Association Studies. PLoS ONE, 2007. 2(7): p. e591.

21. Carlson, J., et al., Widespread Impact of HLA Restriction on Immune Control and Escape Pathways of HIV-1. Journal of Virology, 2012. 86(9): p. 5230–5243.

22. Bhattacharya, T., et al., Founder Effects in the Assessment of HIV Polymorphisms and HLA Allele Associations. Science, 2007. 315(5818): p. 1583–1586.

23. Palmer, D., et al., Integrating genealogical and dynamical modelling to infer escape and reversion rates in HIV epitopes. 2013.

24. Kessinger, T., A. Perelson, and R. Neher, Inferring HIV Escape Rates from Multi-Locus Genotype Data. Frontiers in immunology, 2013. 4.

25. Beerenwinkel, N., et al., Diversity and complexity of HIV-1 drug resistance: A bioinformatics approach to predicting phenotype from genotype. Proceedings of the National Academy of Sciences, 2002. 99(12): p. 8271–8276.

26. Carlson, J., et al., Phylogenetic dependency networks: inferring patterns of CTL escape and codon covariation in HIV-1 Gag. PLoS computational biology, 2008. 4(11): p. e1000225.

27. Carlson, J., et al., HIV-1 adaptation to HLA: a window into virus-host immune interactions. Trends in microbiology, 2015. 23(4): p. 212–224.

28. Chen, L. and C. Lee, Distinguishing HIV-1 drug resistance, accessory, and viral fitness mutations using conditional selection pressure analysis of treated versus untreated patient samples. Biology direct, 2006. 1(1): p. 14.

29. Chen, L., A. Perlina, and C. Lee, Positive selection detection in 40,000 human immunodeficiency virus (HIV) type 1 seguences automatically identifies drug resistance and positive fitness mutations in HIV protease and reverse transcriptase. Journal of virology, 2004. 78(7): p. 3722–3732.

30. Wilson, D. and G. McVean, Estimating diversifying selection and functional constraint in the presence of recombination. Genetics, 2006. 172(3): p. 1411–1425.

31. Nielsen, R. and Z. Yang, Likelihood Models for Detecting Positively Selected Amino Acid Sites and Applications to the HIV-1 Envelope Gene. Genetics, 1998. 148(3): p. 929–936.

32. van Westen, G., et al., Significantly improved HIV inhibitor efficacy prediction employing proteochemometric models generated from antivirogram data. PLoS computational biology, 2013. 9(2).

33. Menéndez-Arias, L., A. Mas, and E. Domingo, Cytotoxic T-lymphocyte responses to HIV-1 reverse transcriptase (review). Viral immunology, 1998. 11(4): p. 167–181.

34. Liu, Y., et al., Selection on the human immunodeficiency virus type 1 proteome following primary infection. Journal of virology, 2006. 80(19): p. 9519–9529.

35. Liu, Y., et al., Dynamics of viral evolution and CTL responses in HIV-1 infection. PloS one, 2011. 6(1).

36. Liu, Y., et al., Evolution of human immunodeficiency virus type 1 cytotoxic T-lymphocyte epitopes: fitness-balanced escape. Journal of virology, 2007. 81(22): p. 12179–12188.

37. Collection of B-list epitopes.

38. Gatanaga, H., et al., Naturally selected rilpivirine-resistant HIV-1 variants by host cellular immunity. Clinical infectious diseases: an official publication of the Infectious Diseases Society of America, 2013. 57(7): p. 1051–1055.

39. Ansari, A., et al., Genome-to-genome analysis highlights the effect of the human innate and adaptive immune systems on the hepatitis C virus. Nature Genetics, 2017. 49(5): p. 666–673.

40. Price, A., et al., Principal components analysis corrects for stratification in genome-wide association studies. Nature Genetics, 2006. 38(8): p. 904–909.

41. Li, N. and M. Stephens, Modeling Linkage Diseguilibrium and Identifying Recombination Hotspots Using Single-Nucleotide Polymorphism Data. Genetics, 2003. 165(4): p. 2213–2233.

42. Tenzer, S., et al., HIV-1 Adaptation to Antigen Processing Results in Population-Level Immune Evasion and Affects Subtype Diversification. Cell Reports, 2014. 7(2): p. 448–463.

43. Llano, A., et al., Best-Characterized HIV-1 CTL Epitopes: The 2013 Update. 2013.

44. Short-Course Antiretroviral Therapy in Primary HIV Infection. N Engl J Med, 2013. 368(3): p. 207–217.

45. Benson, D., et al., GenBank. Nucleic Acids Research, 2014. 42(D1): p. D32–D37.

46. Pan, C., et al., The HIV positive selection mutation database. Nucleic acids research, 2007. 35(Database issue): p. D371–D375.

47. Fagard, C., et al., A prospective trial of structured treatment interruptions in human immunodeficiency virus infection. Archives of internal medicine, 2003. 163(10): p. 12201226.

48. Frater, A.J., et al., Effective T-Cell Responses Select Human Immunodeficiency Virus Mutants and Slow Disease Progression. Journal of Virology, 2007. 81(12): p. 6742–6751.

49. Huang, K.-H., et al., Progression to AIDS in South Africa Is Associated with both Reverting and Compensatory Viral Mutations. PLoS ONE, 2011. 6(4): p. e19018.

50. Huang, K.-H.G., et al., Prevalence of HIV type-1 drug-associated mutations in pre-therapy patients in the Free State, South Africa. Antiviral therapy, 2009. 14(7): p. 975–984.

51. Leslie, A., et al., Additive Contribution of HLA Class I Alleles in the Immune Control of HIV-1 Infection. Journal of Virology, 2010. 84(19): p. 9879–9888.

52. Matthews, P., et al., Central Role of Reverting Mutations in HLA Associations with Human Immunodeficiency Virus Set Point. Journal of Virology, 2008. 82(17): p. 8548–8559.

53. Shapiro, R.L., et al., Antiretroviral Regimens in Pregnancy and Breast-Feeding in Botswana. N Engl J Med, 2010. 362(24): p. 2282–2294.

54. Los Alamos HIV Seguence Database.

